# Membrane rupture and independent extension of sister membranes drive cytokinesis in *C. elegans* embryos

**DOI:** 10.1101/2025.07.26.666980

**Authors:** Jingjing Liang, Tingrui Huang, Xun Huang, Mei Ding

## Abstract

Cell division, the physical separation of a cell into two daughter cells, is classically thought to require continuous plasma membrane invagination driven by equatorial contractile forces. Here, we challenge this paradigm by uncovering an unexpected mechanism in *C. elegans* embryos: the plasma membrane ruptures at metaphase or earlier, generating physically independent sister membranes. This separation of sister membranes persists into later cytokinesis, suggesting that the extension of free-ended membrane could direct cytoplasmic partitioning. The rupture of the mother cell membrane and the independent extension of individual sister membranes create widely distributed membrane discontinuities in early *C. elegans* embryos, yet massive cytoplasmic leakage does not occur. EM observations suggest that exposed cytoplasmic regions are shielded by plasma membranes from neighboring cells or by extracellular matrix components. These findings overturn the long-standing assumption that cytokinesis requires persistent membrane integrity and offer fresh insights into cell division.

## Introduction

The plasma membrane integrity is the corner stone of cell biology. Based upon, current models further propose that cell division should also occur with uninterrupted membrane integrity and the invagination of this intact membrane, driven by a contractile ring of actin filaments, myosin II, and regulatory proteins (Pollard and O’Shaughnessy, 2019), with vesicle fusion supplying new membrane to accommodate shape change (Gerien and Wu, 2018), progressively deepens until pinching the cell into two daughter cells (Figure 1A). Critically, the foundational premise of uninterrupted membrane integrity during cytokinesis has never been rigorously tested until our unexpected discovery in *C. elegans* embryos.

**Figure 1.**
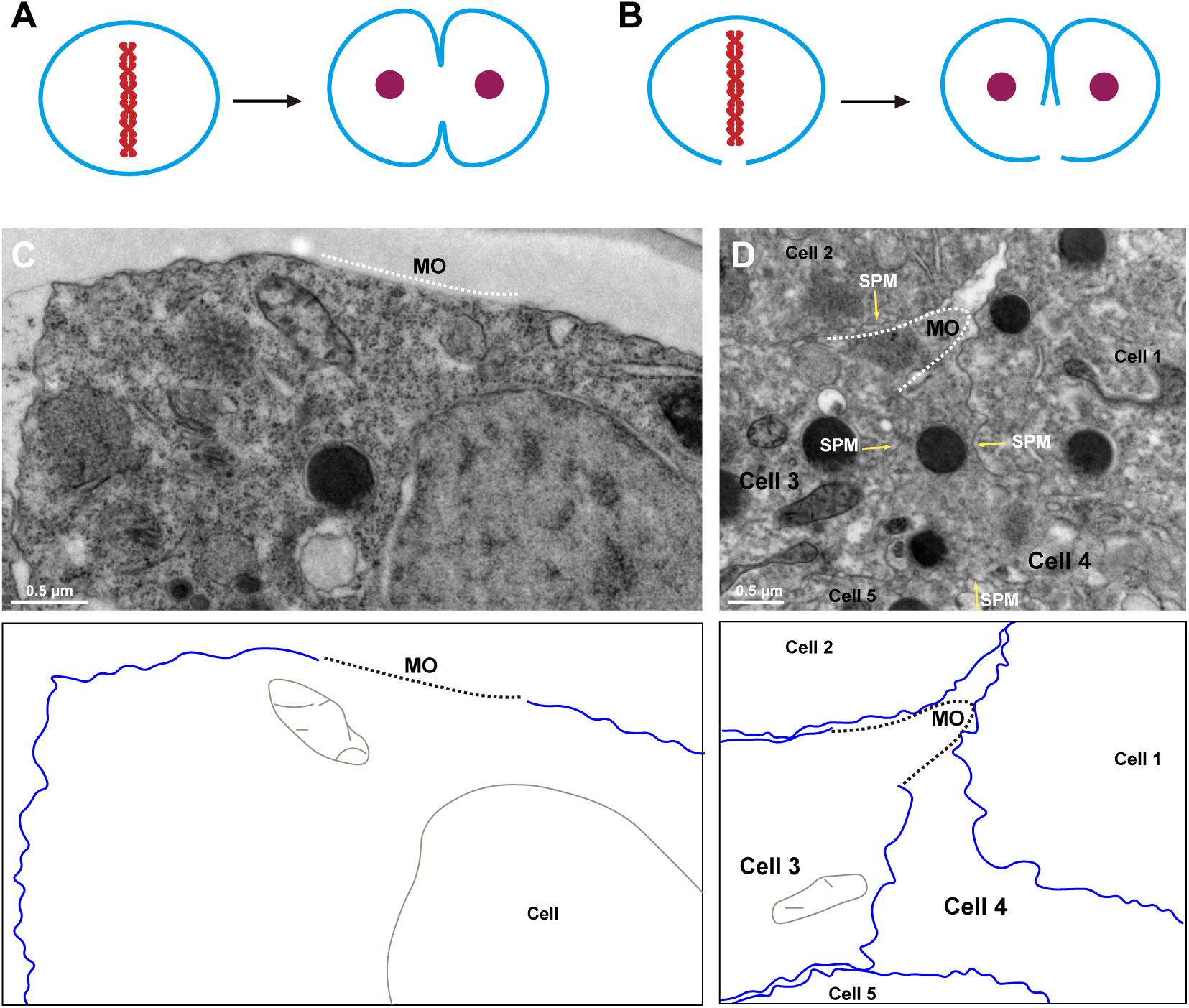
Membrane discontinuities in *C. elegans* embryonic cells. A, Schematic drawing of a metaphase cell and the classical membrane invagination model. B, Schematic drawing of a metaphase cell with a ruptured plasma membrane and a cytokinesis model showing that the detached sister membranes mediate membrane extension in *C. elegans* embryonic cells. C, A membrane opening (MO, indicated by white dashed line) in an embryonic cell at the embryo’s periphery. D, Membrane openings in the embryo’s central regions. Yellow arrows point to single plasma membranes (SPM) traversing the cytoplasm. Plasma membranes are in blue.

After fertilization, the single cell *C. elegans* embryo begins a series of highly stereotyped cell divisions. During this phase, cells undergo rapid division (15-30 minutes per cycle), with each cycle alternating between S (DNA synthesis) and M (mitosis and cytokinesis) phases while bypassing G1and G2 gap phases until the 28-cell stage (Bao et al., 2008; Deppe et al., 1978; Edgar and McGhee, 1988; Pintard and Bowerman, 2019; Sulston et al., 1983). Around this stage, the worm embryos are laid by the mother. Subsequent development continues outside the mother, encapsulated within a protective, multi-layer extracellular matrix known as the eggshell (Stein, 2018). This impermeable eggshell is established earlier, by the end of anaphase II of meiosis, and undergoes its final modifications during the embryo’s first few mitotic divisions (Bembenek et al., 2007; Zhang et al., 2005). Embryos remain within the eggshell and continue to develop until hatch and become young larval worm.

During routine ultrastructural analysis (Zhu et al., 2022), we found that instead of maintaining membrane continuity, dividing *C. elegans* embryonic cells routinely exhibit plasma membrane discontinuities. Further analysis revealed that the plasma membrane ruptures at metaphase or earlier, and the sister membranes primarily extend independently of each other to achieve cytoplasmic partitioning (Figure 1B), a mechanism never previously documented. This discovery challenges the canonical view of cytokinesis at its core, revealing an unanticipated plasticity in cell division mechanisms and prompting a re-evaluation of how membrane dynamics are regulated across species.

## Results

### Membrane discontinuities in *C. elegans* embryonic cells

Our study was prompted by the curious observation that the plasma membrane of embryonic cells in *C. elegans* is not continuously intact. We refer to these membrane discontinuities as membrane openings (MOs). They are particularly evident at the embryo’s periphery: high-pressure freezing transmission electron microscopy (HPF-TEM) revealed that the plasma membrane is absent in the region marked by the white dashed line. Membrane interruptions also occur in central regions. For instance, Figure 1D shows that Cell 3 contains a MO facing both Cell 1 and Cell 2. Additionally, we observed single plasma membranes (SPM) traversing the cytoplasm and appearing to partition it (Figure 1D, yellow arrows). This phenomenon is highly unusual. According to the conventional model, the plasma membrane should remain intact throughout the cell cycle—during both division or interphase—and each cell should be enclosed by its own membrane. Thus, between two adjacent cells, two distinct plasma membranes should be present. Consequently, the observation of a single continuous membrane separating adjacent cells would imply that one of them lacks a complete membrane boundary. Such is the case for Cell 4 in Figure 1D, which clearly exhibits membrane discontinuities, with its exposed cytoplasmic region bounded by membranes from Cells 1, 2, 3 and 5 (Figure 1D).

EM preparation yields serial sections, and the MOs could subsequently be verified with consecutive sections. Two representative examples are displayed in Figure 2. As shown in Figure 2A, three continuous sections capturing one same MO (white dashed lines). On these sections, plasma membranes (marked by yellow arrows) are present on the left and lower sides of the MO. In the enlarged images (Figure 2B and 2C), the junction between membrane and non-membrane regions is visible. If the width of a MO on a given section is calculated based on the direct measurement between two membrane ends, then the MO on these three sections would be approximately 2 μm in width.

**Figure 2.**
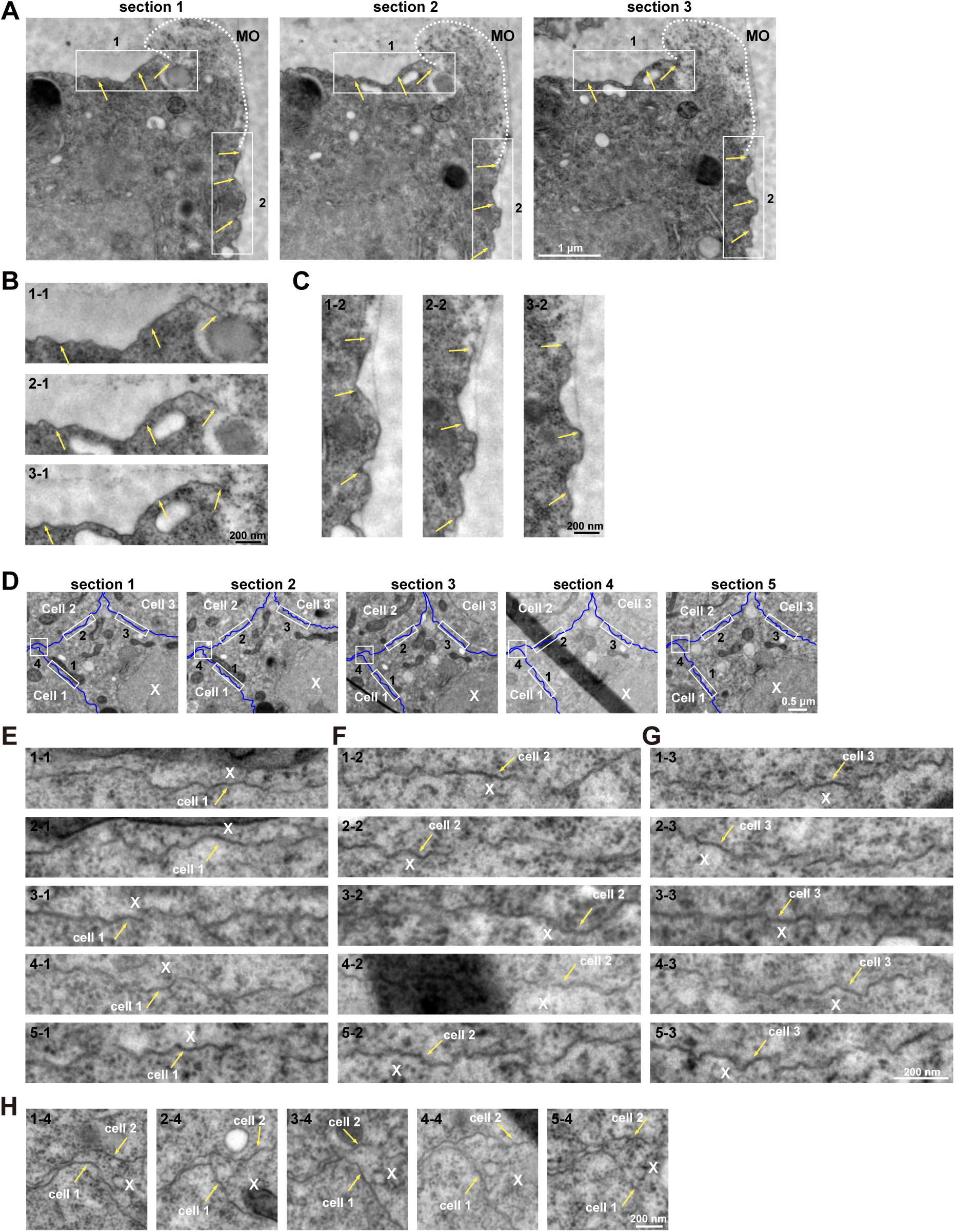
Membrane discontinuities on serial adjacent sections. A-C, The membrane opening (MO) (white dashed lines) through adjacent HPF-EM sections. Plasma membranes are indicated by yellow arrows. The boxed region in (A) is enlarged in (B) and (**C**). D-H, Single plasma membranes traversing the cytoplasm through adjacent HPF-EM sections. Plasma membranes are in blue in (D). Yellow arrows point to single plasma membranes traversing the cytoplasm in (E-H). The boxed region in (D) is enlarged in (E-H).

Single membranes traversing the cytoplasm also indicate the existence of MOs. In Figure 2D, five continuous sections display three single-membrane regions, derived from Cell 1, Cell 2 and Cell 3, respectively (Figure 2E-2G). On these five continuous sections, the cell marked with “X” lacks its own plasma membrane; its exposed cytoplasm is instead shielded by membranes from Cells 1, 2 and 3 (enlarged views of the white rectangular white boxes in Figure 2 are shown in Figure 2E-2G). Figure 2H shows where the Cell 1 and Cell 2 membranes separate (white square), with cytoplasm of Cell X lying between them. Similarly, the cytoplasm of Cell X also lies between the Cell 2 and Cell 3 membranes (Figure 2D). Thus, Cell X exhibits a MO facing Cells 1, 2 and 3.

### The occurrence of membrane discontinuities is developmentally regulated

To rule out potential artifacts caused by EM preparation, we performed additional control experiments. First, we prepared four independent sample batches and consistently observed membrane discontinuities in all them, indicating that this phenomenon is not specific to a single preparation. Second, we examined a publicly available historical collection of *C. elegans* embryo EM images (Wormatlas) and identified MOs in several instances, particularly at the embryonic periphery. This confirmation in an independent dataset suggests that MOs are genuine cellular structures rather than procedural artifacts. Third, we compared intrauterine and already laid embryos and found MO-containing cells in both cases (Figure S1), indicating that the presence of the worm body does not affect the presence of MOs.

In *C. elegans*, embryogenesis begins with rapid cell divisions that quickly increase cell number. Gastrulation (around 28-cell stage) follows, with specific cells moving inward to form the three germ layers. Subsequently, division rates decline as cells adopt specialized fates. The embryo then elongates and develops into a motile larva, which hatches and leaves the eggshell (Figure S2). To understand how MOs arise, we further analyzed embryos across various developmental stages. We found that as embryos reached the comma stage—a period characterized by the onset of elongation and a decline in active cell proliferation—the incidence of MOs dropped dramatically (observed in 0/13, 0/17, 0/30, and 0/30 cells examined, respectively). These results demonstrate that plasma membrane discontinuities are associated with specific stages of *C. elegans* embryogenesis. Moreover, since the eggshell persists throughout embryogenesis, the stage-specific loss of discontinuities cannot be attributed to eggshell-related fixation artifacts.

### Membrane discontinuities are widely distributed in early embryos

Given the specific reduction of MOs in later embryonic stage, we systematically examined how broadly membrane discontinuities are distributed in early *C. elegans* embryos. In total, we collected HPF-TEM images of 60 embryos from 9 gravid adults (Figure S2). By examining the entire circumference of 425 cells in single sections, we identified 159 MO-containing cells (Figure 3A) distributed across 57 embryos, representing 95% of the total sampled embryos. Of note, each membrane discontinuity was identified only after examining the entire plasma membrane of the cell in a given section. Aside from the discontinuity, the remainder of the plasma membrane was intact. Furthermore, all MOs were verified by examining adjacent sections. Among these 159 affected cells, 89% (141/159) exhibited a single membrane discontinuity, while 13 cells had two openings and 5 cells had three.

**Figure 3.**
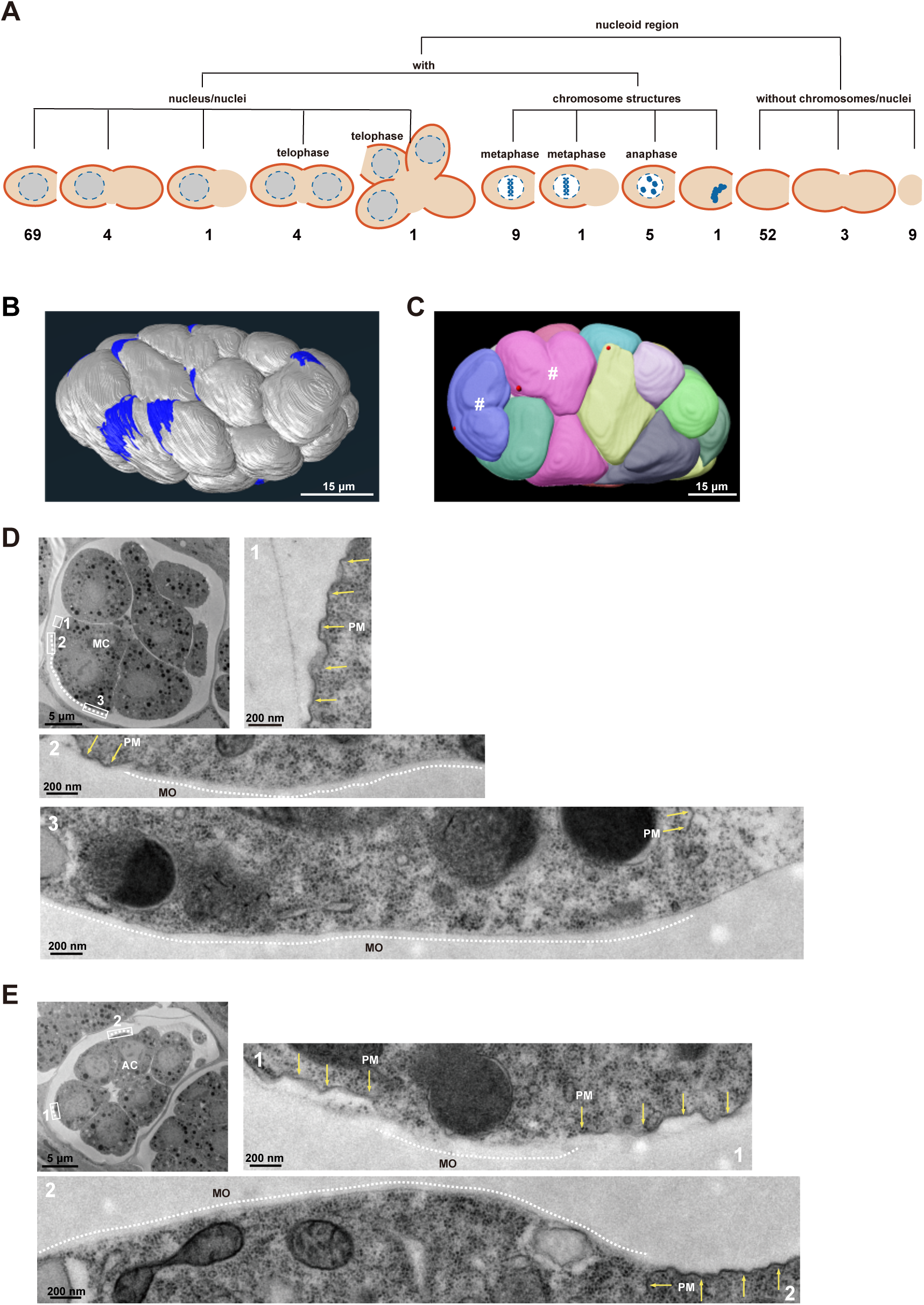
Membrane discontinuities in dividing cells. A, Summary of the single HPF-EM section survey of plasma membrane continuity. Orange lines outline the plasma membranes. Solid gray circles/ellipses enclosed by dashed lines represent evenly stained nucleus regions. Condensed chromosome structures are colored gray-blue. Light brown areas represent cytoplasmic regions. B, Three-dimensional reconstruction of an entire embryo with FIB-SEM. The membrane openings are highlighted in blue. C, Two telophase cells (#), each containing a membrane opening (red dot), were identified in this embryo. D, Membrane discontinuities or openings (MO) (white dashed line) in a metaphase cell (MC). Enlarged views of the boxed regions (1, 2 and 3) are shown. Yellow arrows indicate the plasma membrane (PM). E, MOs (marked by 1 and 2) are detected in anaphase (AC) and other early embryonic cells; enlarged views of the MOs are shown (white dashed lines). All yellow arrows point from the interior of the cell toward the plasma membrane.

To determine the three-dimensional distribution of MOs, we collected 1,093 continuous EM sections using FIB-SEM (focused ion beam scanning electron microscopy), of which 340 were used for three-dimensional reconstruction of an entire embryo (Video S1 and Figure 3B and 3C). This embryo contained 23 nuclei, indicating it was in an early proliferative stage. After examining membrane integrity in every cell across every section, we identified 13 MO, each in a distinct cell. These 13 MOs were distributed across various embryonic cells, and each MO-containing cell exhibited only one MO. Furthermore, consistent with our observations in single EM sections, MOs were present on both the surface and interior of the embryo. Taken together, these results indicate that membrane discontinuities are widely distributed among early embryonic cells in *C. elegans*.

### Membrane discontinuities in cells under division

The early developmental stage of *C. elegans* embryos is characterized by rapid and extensive cell division. Within approximately 100 minutes, a two-cell embryo develops into an embryo containing nearly 30 cells (Bao et al., 2008). Given this rapid proliferation feature of early *C. elegans* embryos, we suspected that the formation of membrane discontinuities might be associated with cell division. To test this possibility, we analyzed the 159 cells exhibiting MOs based on nuclear and cytoplasmic morphology (Figure 3A). Of these, 64 cells appeared in EM sections containing only cytoplasmic regions-9 of which completely lacked discernible plasma membranes. The remaining 95 cells contained one or more nuclei or condensed chromosomes. Among them, 10 cells were clearly in the metaphase stage (defined by a single chromosomal cluster). As shown in Figure 3D, a representative metaphase cell (MC) displayed a MO (dashed white line) spanning roughly one-third of the cell’s circumference. High-magnification imaging revealed that, in contrast to the wrinkled plasma membranes (Figure 3D, enlarged boxed area displayed in 1), the large MO exhibited a smoother surface morphology, and the internal cytoplasm remained enclosed within the cell (Figure 3C, enlarged boxed areas displayed in 2 and 3). MOs were also observed in anaphase cells (AC) (n=5; characterized by <10 chromosomal clumps; Figure 3E, enlarged in 2) and telophase cells (n=5; defined by two nuclei sharing cytoplasm). Intriguingly, one telophase-like cell contained three nuclei within a common cytoplasm (Figure S2, marked by &). Through whole-embryo reconstruction (Video S1), we also identified two telophase cells, each containing a membrane opening (marked by # in Figure 3C). Taken together, these analyses demonstrate that membrane discontinuities are present on cells during mitosis.

To further assess the relationship between mitosis and MOs, we analyzed 20 cells with unambiguous division features, selected based on nuclear morphology—such as aligned chromosomes (metaphase) or discrete clusters (anaphase)—and cytoplasmic sharing, indicated by two or more nuclei within a common cytoplasm (telophase). Strikingly, 19 out of 20 cells (95%) revealed MOs upon tracing the plasma membrane in single EM sections (Figure S3). Notably, cells with unambiguous division features are more likely to be sectioned through or near their maximal diameter. In contrast, the randomly selected sections used for Figure 3A captured cells at variable planes, largely reducing the likelihood of observing MOs. Nevertheless, this near-ubiquitous association (19/20) strongly suggests that membrane discontinuities are an integral feature of cell division in early *C. elegans* embryos.

### Plasma membrane rupture in the metaphase cell

The MOs appear at metaphase, well before nuclear or cytoplasmic division. To understand the origin of MOs, we performed serial EM sectioning to trace the entire plasma membrane of a metaphase cell (Figures 4A and 4B). A total of 112 serial sections were collected, yielding 822 HPF-TEM images. Among these, 105 sections contained this metaphase cell. Through 3D reconstruction, we identified a continuous plasma membrane rupture extending over 8,940 nm, which is more than two-thirds of the cell’s circumference (Figure 4B). Notably, one end of the membrane rupture appeared long and narrow, while the other was wide and exhibited cytoplasmic leakage (Figure 4B and Video S2). Further analysis revealed that the leaked cytoplasm remained confined to a relatively localized region (Video S2). During metaphase, chromosomes align at the equatorial plane, a key region for subsequent segregation. To explore the spatial relationship between membrane rupture and chromosomal organization, we further reconstructed the entire condensed chromosome region. We found that this large-scale membrane rupture largely encircled the chromosomal disc (Figure 4B and Video S2), spatially aligning with the future segregation zone.

**Figure 4.**
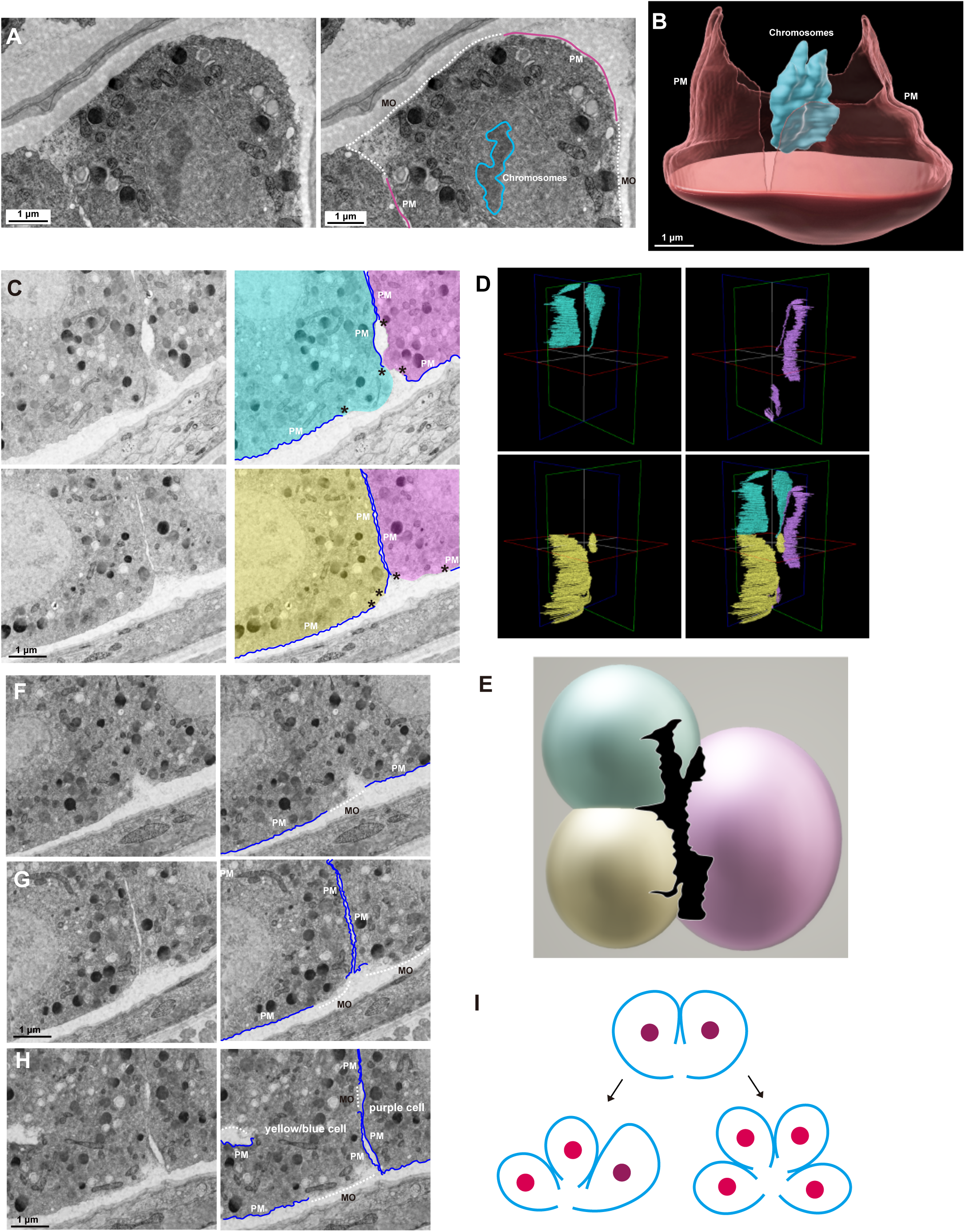
Three-dimensional reconstruction of membrane openings. A and B, Three-dimensional (3D) reconstruction of a metaphase cell. Representative HPF-EM images are shown in (A). Purple lines outline the plasma membranes. White dashed lines outline the exposed cytoplasmic region. Blue lines outline the condensed chromosome region. B, The 3D reconstruction of the metaphase cell. The plasma membrane (PM) and condensed chromosome region are displayed. C, The opposing severed edges are close to each other in the yellow, purple and blue cells. Plasma membranes (PM) are indicated in blue. Asterisks indicate membrane tips. D, Reconstruction of the membrane ruptures and adjacent plasma membranes. Individual membranes, each associated with a distinct nucleus, are color-coded (blue, yellow, and purple). F and G, Membrane openings (MO, dashed white lines) arise either from separation between sister membranes (F) or between the severed membrane edges of the same cell (G). H, In a single EM section, the multiple openings originate from the same rupture. E, The 3D model of a tri-nucleated cell showing the membrane rupture (black). I, Schematic diagram of the formation of cells with three or four nuclei.

Because neither nuclear nor cytoplasmic division is visible during metaphase, the presence of membrane rupture in metaphase cells implies that the sister membranes have already partially separated and the two daughter cells have become at least partially individualized.

### Plasma membrane rupture in a tri-nucleated telophase cell

Membrane rupture in metaphase results in the separation of the two sister membranes. We wondered how the subsequent cytokinesis process proceeds with detached sister membranes. To gain insight into membrane organization during cytokinesis, we applied the AutoCUTS-SEM (automatic collector of ultrathin sections scanning electron microscopy) workflow (Li et al., 2017) to acquire serial EM sections of a telophase cell. Adult worms were treated with high-pressure freezing, freeze-substitution, and embedding as described above. Subsequently, 351 serial sections (60 nm thick) were collected using the AutoCUTS device, and scanning electron microscopy imaging was performed. Through continuous tracing, we discovered that this cell actually contained three independent nuclei. We color-coded each membrane region according to its associated nucleus, with relative sizes estimated from EM sections to ensure proportional representation (Figure 4C). The 3D model demonstrated that the division plane between the yellow and blue cells was roughly perpendicular to that between the purple cell and the other two (Figure 4D).

Using consecutive EM sections, we traced membrane integrity in this tri-nucleated cell. We found that this tri-nucleated cell was enveloped by three distinct plasma membranes, each containing a single rupture site. These three ruptures converged to form a “Y”-shaped exposed cytoplasmic region spanning >351 sections (21,060 nm; Figures 4E and Video S3). Throughout these 351 consecutive sections, we detected no continuity between adjacent membranes, indicating that membrane independence had been established by this stage. Consecutive tracing further showed that the MOs-frequently observed in single EM sections-could arise from either the separation between sister membranes (Figure 4F) or from separated edges of the plasma membrane within the same cell (Figure 4G). Additionally, multiple openings within a single cell might represent different cross-sectional planes of a common membrane rupture (Figure 4H).

In all three cells (where cytoplasmic separation was incomplete), the free membrane edges were in close proximity (marked by *; Figures 4C), indicating that each of these three detached sister plasma membranes had almost completely enclosed its individual cytoplasm. Since membrane rupture can occur in metaphase or even early, this also suggests that detached sister membranes can extend to complete cytokinesis. Moreover, three nuclei sharing a common cytoplasm indicate two concurrent rounds of cytokinesis: the next division began before the previous round finished and even progressed to a later stage of cytokinesis. If membrane rupture occurs at metaphase or earlier, these observations suggest it may persist throughout the cell cycle.

This phenomenon—concurrent divisions—appears to be common in early *C. elegans* embryos, as our EM analysis identified more than 10 cells with three or four nuclei. For example, Sup Figure 4 shows an EM section of four nuclei sharing a common cytoplasm, with MOs identified for Cell 2 and Cell 3 in that section. Based on reconstruction of this tri-nucleated cell, we suspected that a similar mechanism operates in four-nucleated cells, in which two detached daughter membranes-without completing the last round of cell division-undergo further rupture, forming four granddaughter membranes. These detached free-ended membranes then extend independently to complete cell division (Figure 4I).

### The detachment of sister membranes

According to the conventional model, cytokinesis in animal cells begins with membrane invagination, which deepens into two back-to-back membrane layers and ultimately separates two daughter cells. Here, we show that membrane rupture creates separated two sister membranes, and that extension of the free-ended individual sister membrane could mediate cytoplasmic partitioning. To determine how often cytokinesis proceeds with detached sister membranes in early *C. elegans* embryos, we examined 51 dividing cells. To ensure unbiased sampling and consistent comparison, we first identified telophase cells at low magnification (x4,000). At this magnification, all 51 cells displayed two nuclei sharing a common cytoplasm. We then examined the same sections at higher magnification. Again, to ensure fair comparison, all 51 sections were judged at the same magnification (x30,000). Under these consistent conditions, if the sister membranes appeared continuously connected, we defined them as showing “membrane invagination.” In contrast, if the sister membranes appeared separated from each other, we defined them as cells with a “detached membrane configuration.” Using these criteria, we found that a minority of these telophase cells (6/51) appeared to show membrane invagination (Figure 5A), including one embryo transitioning from the one-cell to the two-cell stage. In contrast, the vast majority (45/51) displayed a striking detached membrane configuration (Figure 5B). This predominant pattern suggests that membrane detachment, rather than continuous invagination, may represent the primary mode of cytoplasmic separation in early *C. elegans* embryos.

**Figure 5.**
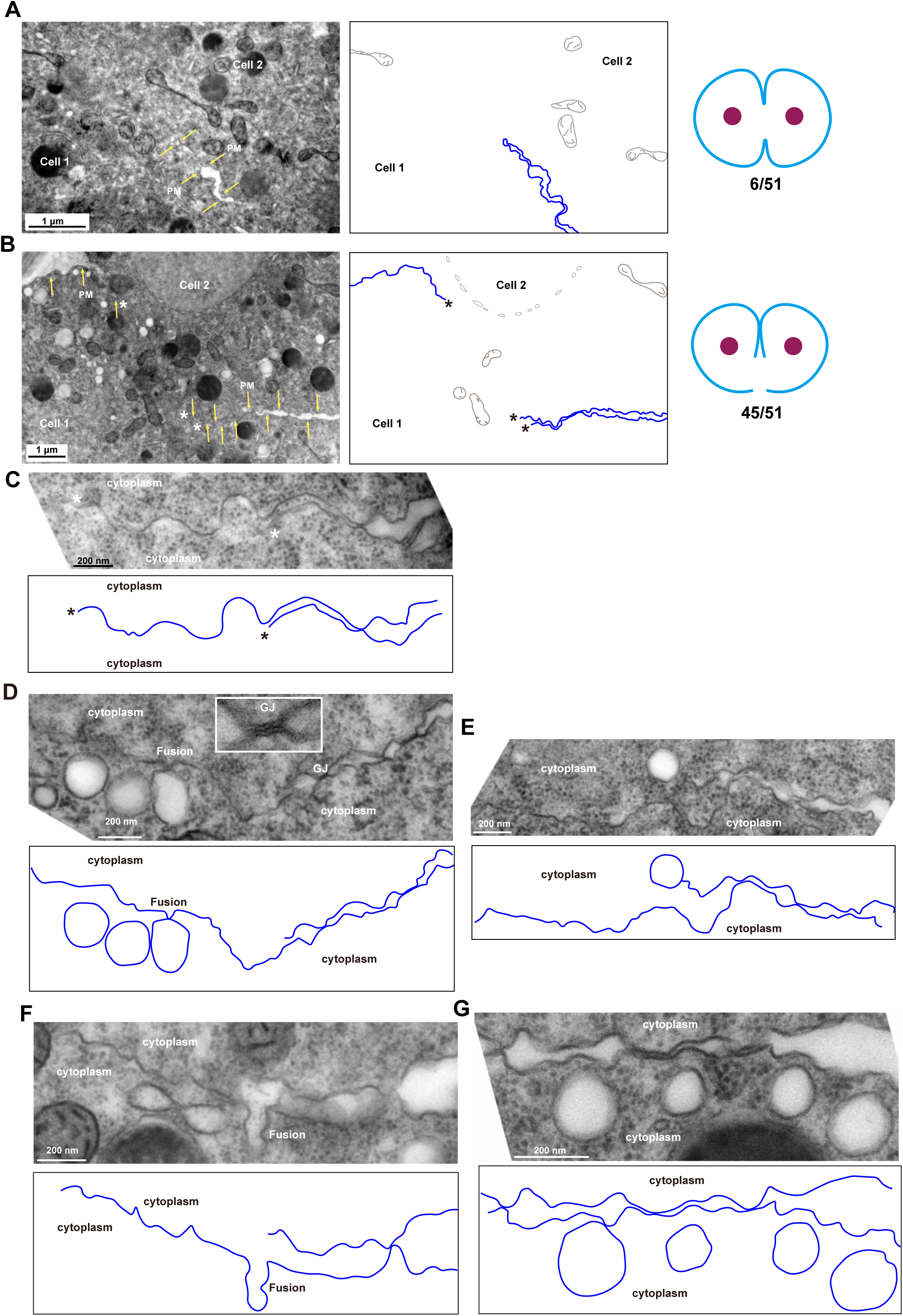
The detached severed sister membranes. A, Representative image of a classical back-to-back membrane arrangement. B, Representative image of detached severed sister membranes. Diagrams of the corresponding cytokinesis models are displayed on the right. C, Tips of two detached sister membranes. D, Vesicles clustering near the cytosol-immersed region of sister membranes. Contact or fusion between vesicles and membranes is indicated. The rectangular inset shows an enlarged view of a gap junction (GJ) plaque. E, A vesicle localized at the tip of a cytosol-immersed membrane. F, Putative fusion events between vesicles and free-ended membranes. G, Vesicles found near the separated sister membranes. Plasma membranes are in blue.

To resolve the detached membrane configuration at high resolution, we acquired HPF-EM images showing the detached edges of the two sister membranes (Figure 5C). In this configuration, the sister membrane tips were completely separated (Figures 5C, marked with *), forming distinct, non-contiguous structures that defy conventional models of cytokinesis. Cytoplasmic components such as glycogen granules and ribosomal particles were clearly visible around the severed edges of the two sister membranes (Figure 5C). This is significant because the classical invagination model cannot generate single plasma membrane barriers between adjacent cytoplasmic domains, whereas independent extension of severed sister membranes could produce cytoplasm-immersed membranes.

We traced 12 such cytoplasm-embedded membranes, including those separating Cell 1 from Cells 2 and 3, and Cell 4 from Cell 3 (Figure S4). In all cases, the cytoplasmic regions flanking these membranes belonged to cytosolically interconnected sister cells, suggesting that the exposed cytoplasmic region may be shielded by membranes from sister cells.

Tracing from the severed edges back to their origins, we found that the sister membranes were frequently anchored side-by-side near the cytosol-immersed region (Figures 5D). Within this region, gap junction plaques were often observed (57 in 94 cases) at membrane contact sites (Figure 5D), though their potential role in sister membrane cohesion is unclear. These contact sites alternated with regions where the two sister membranes separated, revealing lightly stained intercellular spaces in EM images. Beyond the anchored region, the separation between sister membranes became evident and each membrane exhibited the characteristic features of a typical plasma membrane—with one side facing the cytosol and the other exposed to the extracellular environment (Figure 5C and 5D).

Cellular vesicles were observed near free membrane edges (Figures 5D and 5E). Some cytosol-immersed membrane tips appeared connected to vesicles (Figure 5E), with instances of direct contact (Figure 5D) and possible fusion events (Figures 5F). Vesicles also clustered along the anchoring area, as well as the separated sister plasma membranes (Figure 5G). EM analysis revealed no morphological differences between vesicles associated with the cytosol-immersed region and those near conventional regions of the sister membranes.

### Capturing the membrane rupture and independent extension of detached sister membranes in living embryos

While high resolution HPF-TEM analysis revealed widespread membrane discontinuities in early *C. elegans* embryos, this static imaging approach cannot capture dynamic membrane behavior. Next, we employed a fluorescence-labeled membrane marker, GFP::PH(PLC1δ1), to investigate the dynamic movements of membrane ruptures and detached sister membranes. Our live imaging detected both putative membrane ruptures (Figure 6A) and free-ended sister membrane structures (Figures 6B and 6C), providing additional evidence that membrane rupture and independent extension of detached sister membranes underlie cytokinesis in *C. elegans* embryos.

**Figure 6.**
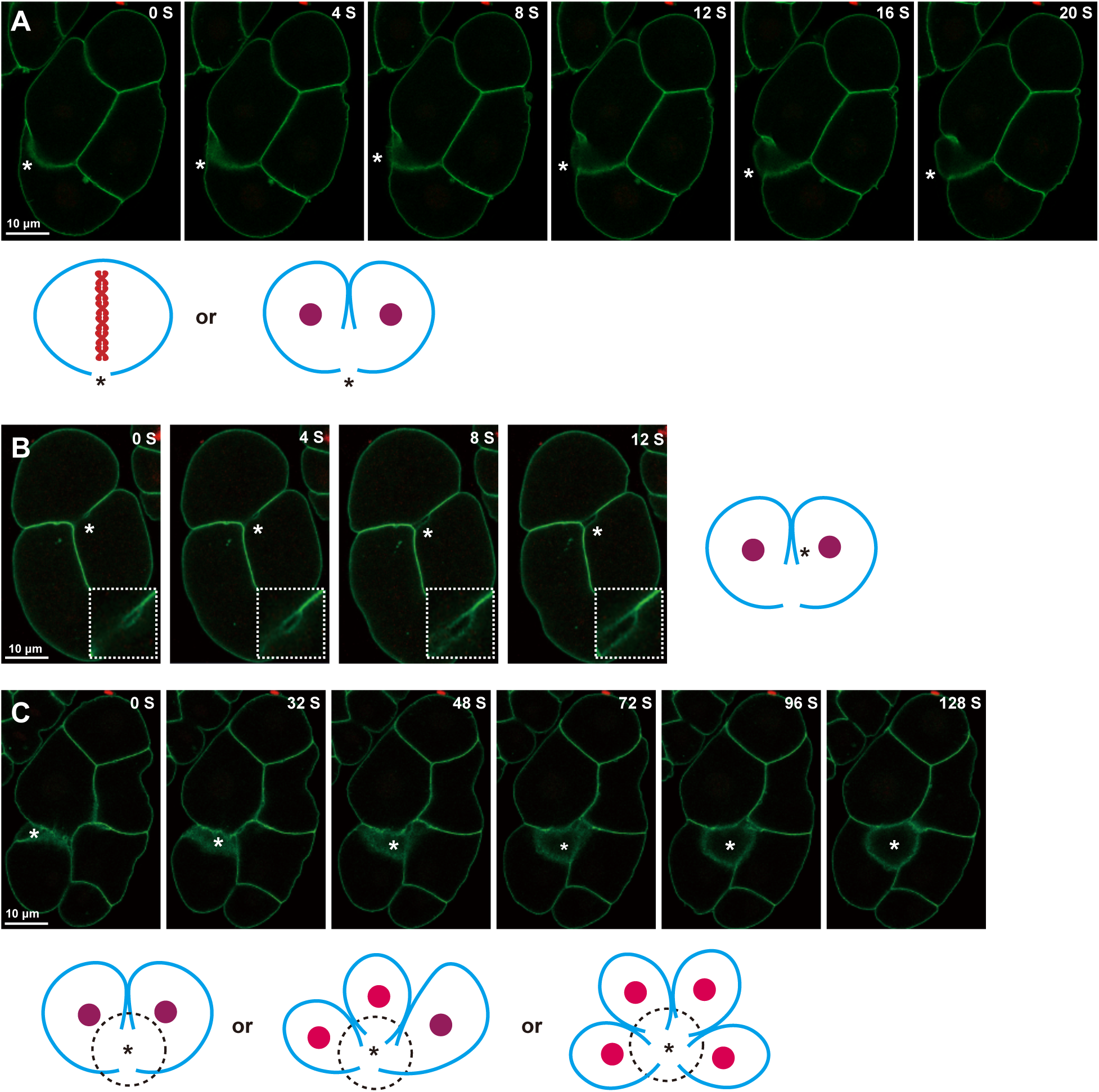
Detection of membrane disruptions by optical microscopy. A, The putative membrane rupture and extension (*). B, Two extending free-ended sister membranes (*). Insets are enlarged views. C, Possible free-ended edges of detached membranes, probably from multiple cells (*). Below are proposed models illustrating the membrane disruptions that could explain the anomalous GFP signals. Images were taken at 4-second intervals for (A and B) and 8-second intervals for (C). Green signal is GFP::PH(PLC1δ1).

To capture the 3-dimensional membrane dynamics, we turned to lattice light-sheet microscopy. Imaging one embryo at a time from the 2- to 4-cell stage, we acquired volumetric stacks spanning the entire embryo in approximately 10-20 seconds, at 20-second intervals. Using the GFP::PH marker, we captured the dynamic movement of large membrane ruptures in early embryos (Video S4 and Figure7A). The appearance and disappearance of these membrane openings provided compelling evidence that membrane rupture is a general feature of early embryonic cells in *C. elegans*.

To reveal the dynamic association between nuclear division and membrane rupture, we further co-labeled chromatin with mCherry::H2B. We found that membrane rupture was present almost throughout the entire mitotic process, including prophase, metaphase, anaphase and telophase (Video S4 and Figure 7B). As the future sister nuclei moved further apart, the membrane opening appeared to break into two openings (Figure 7B). Together with the EM observations, this suggests that the individual detached sister membranes were extending to complete the enclosure of the cytoplasm.

**Figure 7.**
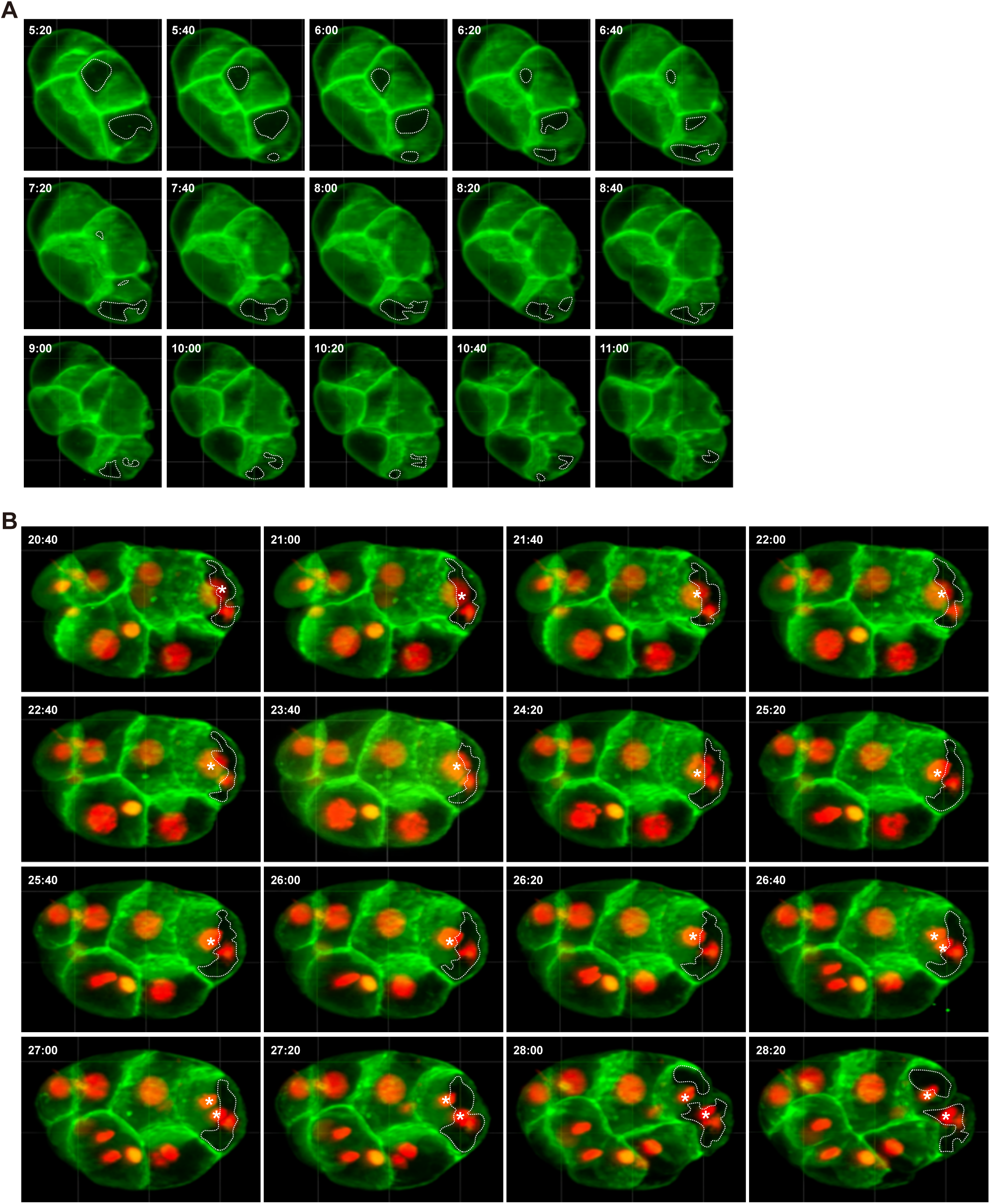
Dynamics of three-dimensional membrane openings visualized by lattice light-sheet microscopy. A, Dynamic movement of membrane openings in early embryos. B, Dynamic movements of nuclear division and membrane opening. Green signal is GFP::PH(PLC1δ1). Red signal is mCherry::H2B. Asterisks mark the nucleus undergoing division.

### Sealing the exposed cytosol by neighboring membranes and extracellular matrix layers

Conventionally, the integrity of the plasma membrane is considered essential for cell survival, and its disruption can lead to cytoplasmic leakage and, in severe cases, cell death. However, widespread cytoplasmic leakage has not been observed in wild-type *C. elegans* embryos. Based on our EM observations, we propose two mechanisms through which dividing embryonic cells may seal or cover exposed cytosolic regions.

First, the sharing of plasma membrane with neighboring cells may protect the exposed cytosol. As repeatedly shown in EM images in this study (Figure 1D, 2D–2H, 5C and S4), a single membrane shields not only its own cytoplasm but also that of neighboring cells. These neighboring cells may be direct sister cells derived from the same mother cell, or cousin cells originating from mother cells that were once sisters. In cases where three nuclei share a common cytosol, the adjacent cell could even be an aunt or nephew cell.

In addition, eggshell components may seal cytoplasmic regions lacking membrane coverage. The *C. elegans* eggshell is composed of six distinct extracellular matrix layers (Olson et al., 2012; Benenati et al., 2009; Rappleye et al., 1999; Schierenberg and Junkersdorf, 1992; Stein, 2018). Among these, the fifth permeability barrier layer (PBL), a lipid-rich structure, serves as the osmotic and permeability barrier for the developing embryo (Benenati et al., 2009; Olson et al., 2012). Normally, the PBL encloses the peri-embryonic layer (PEL), which separates it from the embryonic plasma membrane (Benenati et al., 2009; Olson et al., 2012) (Figures 8A and 8B). In contrast, in regions where the plasma membrane was absent, we observed the PBL adhering directly to the exposed cytoplasm (red triangles and dashed red lines, Figures 8B and 8C), bypassing the PEL. This phenomenon was also evident in other EM images (Figures 2C, 3C, S1 and S3). PBL-mediated sealing is not always complete. When the PBL was not accessible or only loosely covered membrane-free regions, minor cytoplasmic leakage occurred (Figure 8D). However, the leaked cytoplasmic material appeared to remain confined region, likely by the PEL (Figure 8D). The mechanisms by which extracellular matrix components protect the exposed cytoplasmic region are unknown.

**Figure 8.**
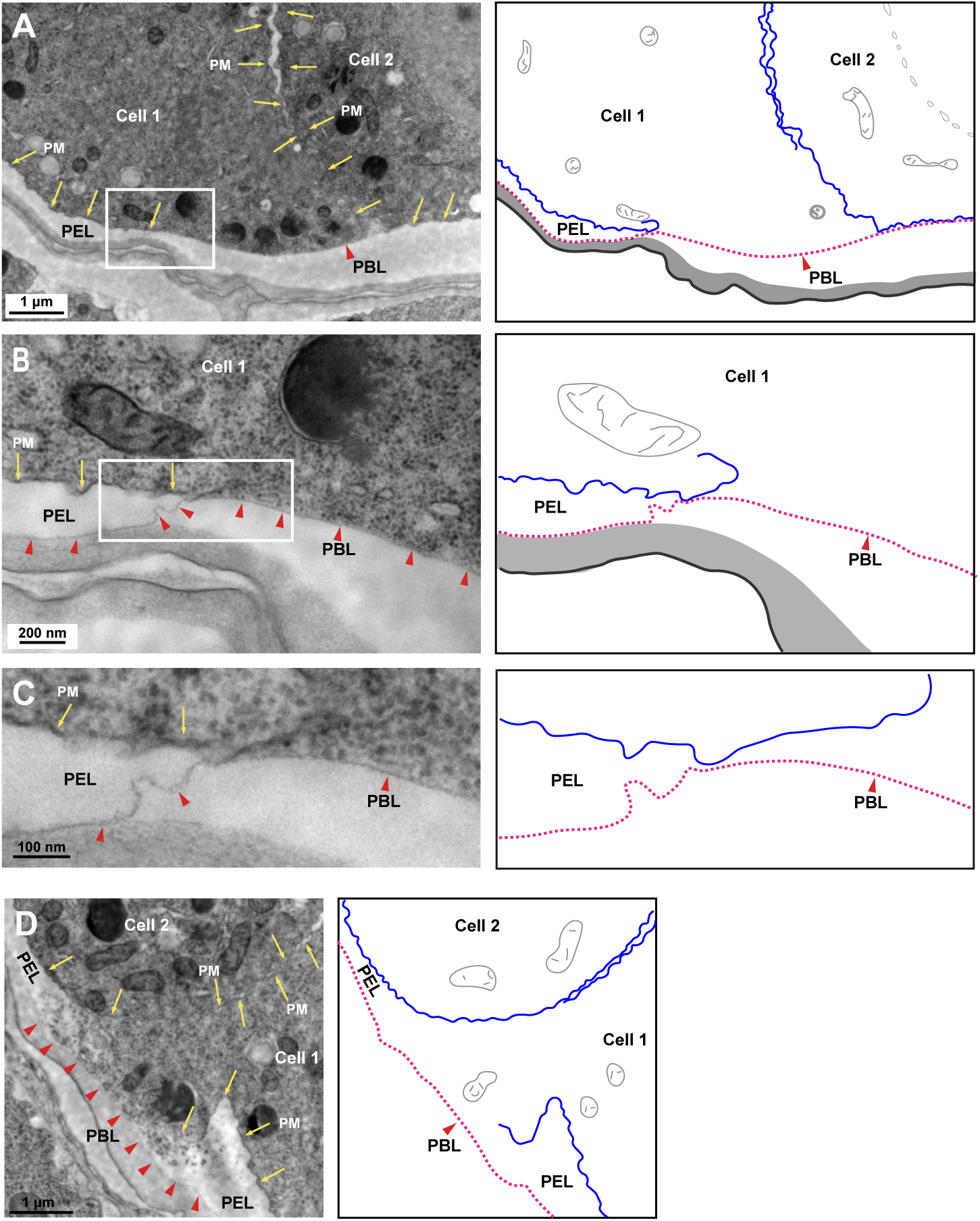
The exposed cytoplasmic regions covered by extracellular matrix layers. A-C, The exposed cytoplasmic region is covered by the permeability barrier layer (PBL, red triangles and red dashed lines). PM, plasma membrane (yellow arrows and blue lines). The boxed region in (A) is enlarged in (B), and the boxed region in (B) is enlarged in (C). D, The exposed cytoplasmic region can also be confined by the peri-embryonic layer (PEL).

## Discussion

In this study, we discovered that the plasma membrane of *C. elegans* embryonic cells undergo rupture during cell division. Following this rupture, the detached sister membranes extend independently from each other to complete cytokinesis. To date, the ultrastructural dynamics of plasma membrane integrity during cell division remained unexplored in other species. We anticipate that our findings will catalyze comparative studies across different species or under various pathological conditions, to determine whether this mechanism represents a fundamental paradigm or a context-specific strategy. Our work established the first mechanistic framework to guide such future investigations.

### Partitioning cytoplasm with detached sister membranes

Cell division involves both nuclear and cytoplasmic division. However, due to the difficulty of visualizing cell boundaries in living embryos, previous studies in *C. elegans* have largely emphasized nuclear events. Here, we observed that early *C. elegans* embryonic cells can contain three or four nuclei within a shared cytoplasm, indicating that cytoplasmic division may be uncoupled from nuclear division. This phenomenon is not unique to *C. elegans*—it has also been reported in other species such as *Drosophila*, where the early embryo undergoes 13 rapid nuclear divisions without cytokinesis before cellularization occurs. It has been proposed that by minimizing or delaying cytokinesis, embryos can accelerate nuclear multiplication, thereby supporting the rapid pace of early development. In this context, the embryonic cells of *C. elegans* appear to adopt a notably rapid—though seemingly risky—strategy for cytoplasmic partitioning, yet one with clear developmental benefits.

First, membrane rupture may help alleviate the rise in intracellular pressure during division (Hiramoto, 1963; Stewart et al., 2011). Rupture occurs at metaphase or earlier and can persist throughout the cell cycle. An opening in the membrane likely reduces internal pressure, thereby conserving energy and resources that would otherwise be expended on resisting membrane tension. These conserved resources could then be redirected toward essential processes such as chromosome segregation and membrane synthesis-a particular advantage for rapidly dividing embryonic cells. Notably, membrane rupture occurs at metaphase or even earlier, before nuclear or cytoplasmic division, suggesting that the physical independence of the plasma membranes may serve as an early marker of nascent daughter cells, preceding other cellular landmarks.

Following membrane rupture, the sister membranes become separated, raising the question of how cytokinesis proceeds with severed membranes. Analysis of single EM sections revealed continuity between sister membranes resembling invagination in only a few cases (6 out of 51), suggesting that while a subset of embryonic cells may employ the membrane invagination and extension model, the majority do not. Interestingly, invagination was detected during the one-cell to two-cell transition, yet membrane discontinuities were observed at this same stage (Figure S2). This indicates that free-ended-membrane-mediated extension may co-occur with invagination. Alternatively, the back-to-back invagination may be transient, subsequently breaking up into free-ended membranes that continue their extension. Distinguishing between these hypotheses remains challenging at present. A definitive resolution will likely require advanced imaging techniques capable of high-resolution, real-time visualization at the EM level.

Unlike a continuous bilayer, a free-ended membrane may offer greater flexibility during cytokinesis. Without being physically constrained by its sister, each membrane can extend in different directions as needed. This independence may be particularly advantageous in accommodating asynchrony between nuclear and cytoplasmic division, as seen in early *C. elegans* embryos where multinucleated cells form through incomplete cytokinesis. In such cases, independent membrane growth from two directions facilitates cytoplasmic separation more effectively than a single continuous membrane would allow. Such autonomous movement may also explain the divergent orientations of sister membranes frequently observed in EM sections.

Through EM analysis, we observed vesicles randomly distributed beneath the extending sister membranes, including at free-ended membrane edges, the side-by-side anchoring area, and the separated sister plasma membranes (Figure 5). EM analysis further showed no morphological differences between vesicles near cytosol-immersed regions and those at conventional membrane sites. Although vesicle fusion is considered an important contributor to sister membrane extension in the conventional model, whether it also plays a role in the extension of free-ended sister membranes remains completely unknown. Further investigations will be needed to test this hypothesis.

Finally, when the separated membrane edges meet within daughter cells, full cytoplasmic separation is achieved. In the classical invagination-extension model, cytokinesis requires both membrane scission and fusion. In the detached-membrane mode described here, however, cytokinesis proceeds without a continuous membrane structure. This raises important questions: Is membrane scission dispensable in this context? Could bypassing scission accelerate cell separation? If so, might such a mechanism offer a selective advantage during rapid embryonic development? These questions remain open and merit further investigation.

### Limitations and perspectives

In this study, the application of HPF-TEM, FIB-SEM and live imaging has enabled unprecedented ultrastructural visualization of dividing cell surfaces in early *C. elegans* embryos. Our findings reveal a cytoplasmic division mechanism that challenges current models and raises important new questions regarding the plasticity of cytokinesis.

HPF-TEM offers a significant advantage for membrane studies by capturing cellular structures in a near-native state. Through ultra-rapid freezing under high pressure, it suppresses ice crystal formation and preserves water in a vitreous state, thereby minimizing chemical artifacts and maintaining the integrity of lipids and proteins. This method provides a faithful snapshot of membrane architecture and dynamics (Moor et al., 1980). In our experiments, the consistent observation of limited number of membrane discontinuities—generally one per dividing cell—that are closely associated with cell division strongly suggests these are genuine biological structures rather than preparation-induced artifacts. While one might argue that such features could reflect EM-related artifacts specific to the plasma membrane during division, the correlative evidence presented here supports their biological relevance.

Using fluorescence-labeled membrane markers, we identified membrane ruptures and free-ended sister membrane structures in live embryos (Figure 6). Light sheet microscopy further resolved the membrane openings and their spatiotemporal relationship with nuclear division and membrane (Figure 7). Intriguingly, similar “abnormal” or “fuzzy” membrane signals have been reported before (Fu et al., 2016; Harrell and Goldstein, 2011), yet the membrane rupture and separated sister membranes failed to be identified. Why? We propose several reasons. First, membrane rupture and free-ended membrane extension are rapid—about 10-30 seconds during metaphase and less than 3 minutes during cytokinesis—and are coupled with cell motility, creating inherent spatiotemporal ambiguities. Consequently, such “fuzzy” or “abnormal” signals were readily dismissed as technical artifacts, whether arising from GFP-based live imaging or EM. More fundamentally, the longstanding dominance of the membrane invagination model has shaped-and potentially constrained-interpretations of membrane dynamics. This paradigm has led researchers to prioritize observations consistent with invagination while dismissing ambiguous structures, such as free-ended extensions, as artifacts rather than considering them as potential elements of alternative mechanisms. Furthermore, theoretical and computational models of cytokinesis have largely been built on invagination-centric assumptions, further narrowing the conceptual framework.

Our findings therefore not only propose a transformative model for cytokinesis but also open a new field of inquiry. Fully validating this mechanism and exploring its conservation and implications will require an integrated approach combining advanced live-cell imaging, biophysical modeling, and genetic tools.

## Methods and Materials

### Examination of *C. elegans* embryonic cells

Wild-type N2 worms were cultured on OP50-seeded NGM plates at 22°C (Brenner, 1974). To obtain images of early embryos, gravid hermaphrodites were collected and sectioned. At low magnification (x4000), individual embryonic cells were identified by the presence of a nucleus. In sections where the nucleus was not visible, cytoplasmic regions—whether covered or not covered by plasma membrane—were counted as individual cells if they were separated from surrounding cells. A total of 425 cells were examined (summarized in Figure 3A), with one EM section analyzed per cell. For embryos with fewer than 10 cells (one or two rows of cells), every cell in the given section was examined. For embryos with three or more rows of cells, only cells at the periphery were analyzed. At higher magnification (x30,000), the entire surface of each cell was examined, revealing 159 cells with one or more membrane openings. Higher magnification improved the resolution of cytoplasmic continuity, reducing potential misidentification of multinucleated cells or interconnected cytoplasmic regions that might have been miscounted at lower magnification. A total of 60 early embryos were serially sectioned as completely as possible, with each embryo yielding at least 20 sections. While Figure 3A presents summary data, images from additional sections were also used for further analyses. Since these additional sections may capture different cells, the total number of embryonic cells examined likely exceeds 425.

Additionally, we examined eight embryos laid outside the worm. Among these, four pre-comma stage embryos each contained at least one MO-positive cell. In another pre-comma stage embryo, no MOs were detected after examining 17 cells. Analysis of one comma-stage embryo (19 cells examined) and two 1.5-fold-stage embryos (>30 cells analyzed each) revealed no MOs.

To determine how frequently cytokinesis proceeds with detached sister membranes in early *C. elegans* embryos, 51 telophase cells were selected on the basis of two nuclei sharing a common cytoplasm at x4,000 magnification. The same sections were then examined at x30,000 magnification. If the sister membranes appeared continuously connected, they were defined as showing “membrane invagination.” If the sister membranes appeared separated from each other, they were defined as showing a “detached membrane configuration.” An additional set of images was taken at x60,000 or x80,000 magnification to reveal the detached sister membranes, and those detached sister membranes were traced using serial sections to confirm that the two membrane tips belonged to sister cells, as judged by these two membranes enclosing a common cytoplasm and each membrane having its own nucleus.

### Identification of MOs

The entire cell surface of individual embryonic cells was examined using electron microscopy at 30,000x magnification. The isolated cytoplasmic regions without plasma membrane coverage were defined as MOs. Sites where the plasma membrane terminated and reappeared were marked as putative MOs. In periphery region of worm embryos, in the non-plasma-membrane-covering regions, the PEL was generally absent, while the PBL typically extended over the exposed cytoplasmic surface. Plasma membranes often appeared wrinkled, whereas cytoplasmic regions covered by the PBL exhibited a smoother appearance. Smooth, non-plasma-membrane-covered regions overlaid by the PBL are readily identifiable as MOs. When exposed cytoplasm was not tightly covered by the PBL, cytoplasmic leakage-still confined to a specific area by the PEL-could also indicate a MO. Given the variable morphology and frequent cytoplasmic leakage, membrane discontinuity size could not serve as a reliable criterion for defining MOs. However, to ensure robustness in our analysis, we considered only linear membrane discontinuities exceeding 200 nm as genuine openings. Each MO was examined independently at least three times. When necessary, adjacent sections were also reviewed to conform an MO’s presence, and identification was finalized only after all three assessments were positive.

### High-pressure freezing and transmission electron microscopy

Basic protocols for high-pressure freezing of a range of organisms, including *C. elegans*, have been described (Manning and Richmond, 2015; McDonald, 2007; Weimer, 2006). Briefly, adult worms were loaded into carriers and cryofixed using a Leica Microsystems EM ICE high-pressure freezer, followed by automatic cooling in liquid nitrogen. The carriers used for freezing (Leica Microsystems, Germany; catalog nos. 16770153 and 16770152) were coated with a non-stick layer of 1-hexadecene (Sigma-Aldrich). After high-pressure freezing (HPF), the samples were transferred under liquid nitrogen to a Leica AFS-2 freeze-substitution unit and incubated at −90°C for 72 h in a freeze-substitution solution consisting of acetone with 2% (wt/vol) osmium tetroxide. The temperature was then raised at a rate of 8°C/h for 4 h, held at −60°C for 12 h, increased again at 5°C/h for 6 h, held at −30°C for 10 h, and finally raised to 10°C at a rate of 4°C/h for 10 h. Samples were washed four times in acetone, stained with 1% uranyl acetate for 1 h, and rinsed three times with pure acetone. They were then infiltrated stepwise in increasing concentrations of SPI-Chem™ resin (2:1 resin:acetone for 3 h, 1:1 for 5 h), followed by two incubations in 100% fresh resin (4 h each). The samples were placed in fresh resin in embedding molds and polymerized in a 60°C oven for 3 days. Ultrathin sections (60 nm) were cut using a diamond knife (Diatome) on an ultramicrotome (Leica Ultracut UC7) and collected on slot copper grids (EMS). The sections were visualized using a Hitachi HT7700 transmission electron microscope operating at 80 kV. Images were recorded with a Gatan 832 (4k x 2.7k) CCD camera. Data from four independent preparations were analyzed in this study.

### 3D reconstruction

The metaphase 3D model was prepared using the following steps. First, resin blocks containing gravid adults were prepared via high-pressure freezing and freeze substitution. Continuous ultrathin sections (60 nm thick) were manually collected using a Leica UC-7 ultramicrotome and mounted on slot grids for TEM imaging. A total of 112 serial sections were collected, though 3 sections were lost due to obstruction, resulting in 822 TEM images. Among these, 105 sections contained metaphase-stage cell information. To examine the plasma membrane integrity, high-magnification imaging was performed. Since a single high-magnification image could not cover the entire cell, multiple overlapping images were captured per section and stitched together using ImageJ to generate a complete composite image of the cellular morphology at each section. In total, 292 images were stitched. The 105 stitched images were then aligned using ImageJ, and 99 of these aligned images were used for 3D reconstruction using Imaris software. About a quarter of the cell lacked membrane discontinuities. For this portion, we estimated its size and shape based on EM sections to ensure proportional representation. Of note, single MOs smaller than 200 nm that appeared continuously across more than 3 consecutive sections were considered the endpoint of the membrane rupture. Based on this, the actual length of the membrane rupture could be underestimated at one end, while the other end was clearly defined.

The ultrastructural 3D study of the tri-nucleated cell was performed using an AutoCUTS device (Li et al., 2017) and a Helios Nanolab 600i dual-beam scanning electron microscope (FEI). Adult worms were treated with high-pressure freezing, freeze-substitution, and embedding as described above. Subsequently, 351 serial sections (60 nm thick) were collected using the AutoCUTS device, and scanning electron microscopy imaging was performed. All datasets were analyzed using IMARIS 8 software. After Z-stacks were aligned, the membrane rupture was manually traced and reconstructed.

### FIB-SEM experimental procedure

Nematode samples were prepared into resin blocks using high-pressure freezing and freeze substitution. Initial ultrathin sectioning was t performed for coarse positioning, followed by X-ray microscope (model: Xradia 610) for accurate three-dimensional positioning of the target sample. Based on the X-ray reconstruction, further ultrathin sectioning brought the target as close as possible to the resin surface to obtain higher-quality images. Finally, serial sectioning data were acquired using a focused ion beam scanning electron micros FIB-SEM (instrument models: Crossbeam 340 & VCT500). A total of 1,093 embryo section images were acquired by FIB-SEM, of which 340 were used for three-dimensional reconstruction of an entire embryo. The XYZ pixel dimensions were 65.66/65.66/100 nm. All data underwent three rounds of alignment using FIGI software, followed by three-dimensional reconstruction in Amira software to yield the final three-dimensional image.

### Live imaging of plasma membrane dynamics using GFP::PH

The plasma membrane was labeled using GFP::PH(PLC1δ1). The strain carrying this marker was obtained from the Caenorhabditis Genetics Center and was originally generated in Dr. Karen Faye Oegema’s laboratory. Gravid adult hermaphrodites expressing this marker were collected and washed with M9 buffer. Embryos were then released from the gravid adult worms using a 1 mL syringe needle and transferred onto a 2% agarose pad. Fluorescence imaging was performed on a Zeiss LSM-980 microscope with a 63x/1.42 oil objective. Only one embryo at a time was imaged, starting at the 2- to 4-cell stage. Images were acquired at 4 or 8-second intervals, with total imaging duration typically kept under 30 minutes to ensure embryo viability.

### Lattice light-sheet microscopy

The plasma membrane was labeled with GFP::PH(PLC1δ1) and chromatin with mCherry::H2B. Gravid adult hermaphrodites expressing either GFP::PH alone or both GFP::PH and mCherry::H2B were collected, washed in M9 buffer, and dissected with a 1 mL syringe needle to release embryos onto a 3% agarose pad. Embryos were covered with a coverslip, and the space beneath the coverslip was filled with M9 buffer. Coverslip edges were sealed with paraffin wax.

Fluorescence imaging was performed on a ZEISS Lattice Lightsheet 7 microscope with a 40x/1.0 water-immersion objective, using the 100×1,800 light-sheet setting for single-labeled GFP::PH imaging and the 30×1,000 setting for dual-label imaging. Images were acquired at 2,048×2,048 pixels with a 0.5 μm step size along the X-axis. A single embryo was imaged at a time, starting from the 2- to 4-cell stage. Acquiring a volumetric stack spanning the entire embryo required approximately 10-20 seconds, and stacks were collected at 20-second intervals. Following acquisition, image stacks were deskewed into conventional z-stacks with a final z-spacing of 0.145 μm. Both single- and dual-label datasets were visualized in three dimensions using ZEISS arivis software (version 4.1.1), and movies were generated from these datasets.

## Acknowledgements

We thank Dr. Yanbao Tian from the Core Facility of State Key Laboratory of Seed Innovation and the Center for Biological Imaging at the Institute of Biophysics, Chinese Academy of Sciences (CAS) for technical support. We are grateful to Drs. John White, Andrew Chisholm, and Zhuo Du for their encouragement and insightful suggestions. This work was supported by grants from the National Key R&D Program of China (2024YFA1803401, 2021YFA0805802, and 2024YFA1306100) and the Natural Science Foundation of China (32170790 and 32321004).

## Author contributions

Study concept and design: MD, JL. EM analyses: JL, TH, MD. Drafting and major editing of original manuscript: MD, JL, TH, XH. All authors reviewed, revised, and approved the final version of the paper.

## Supplementary Figure Legends

**Figure S1.**
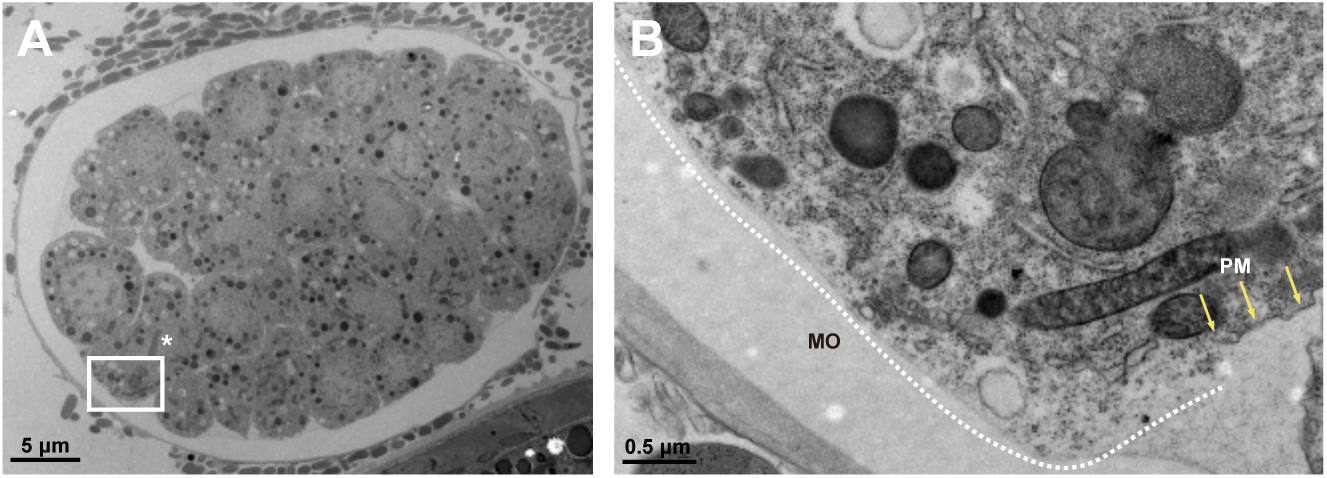
Membrane discontinuities in externally laid *C. elegans* embryos. A, An externally laid embryos with a membrane discontinuity (white box). B, Higher-magnification view of the membrane opening (MO) (white dashed line). Yellow arrow, plasma membrane; asterisk (*), condensed chromosome.

**Figure S2.**
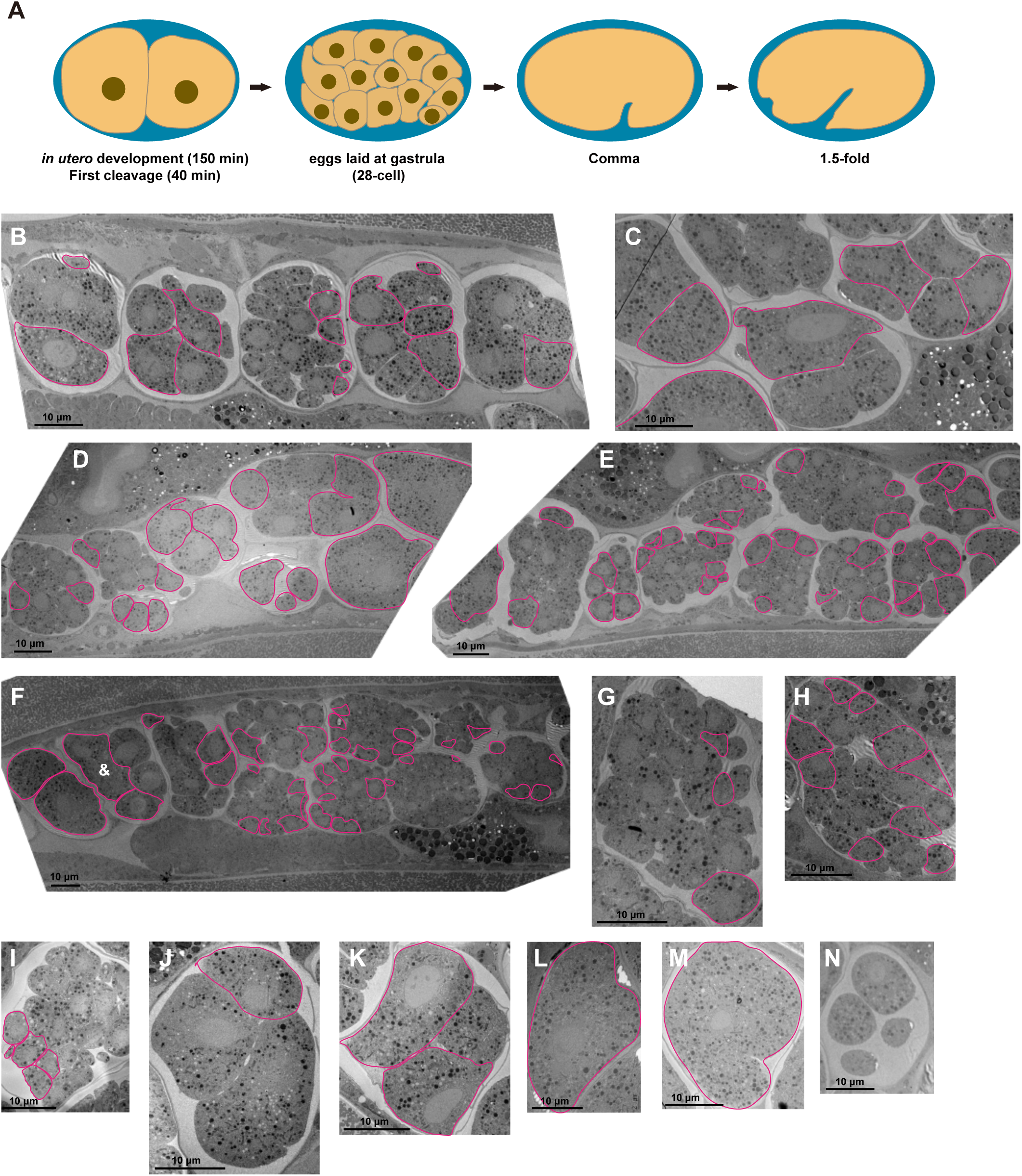
The wide distribution of cells with membrane discontinuities in early *C. elegans* embryos. A, Schematic diagram of *C. elegans* embryogenesis. B-N, Representative low-magnification EM images of early embryos within adult hermaphrodites. Red lines outline cells contain membrane openings. In (F), a tri-nucleated cell is marked with &. (L) and (M) show embryos during the transition from the one-cell to the two-cell stage.

**Figure S3.**
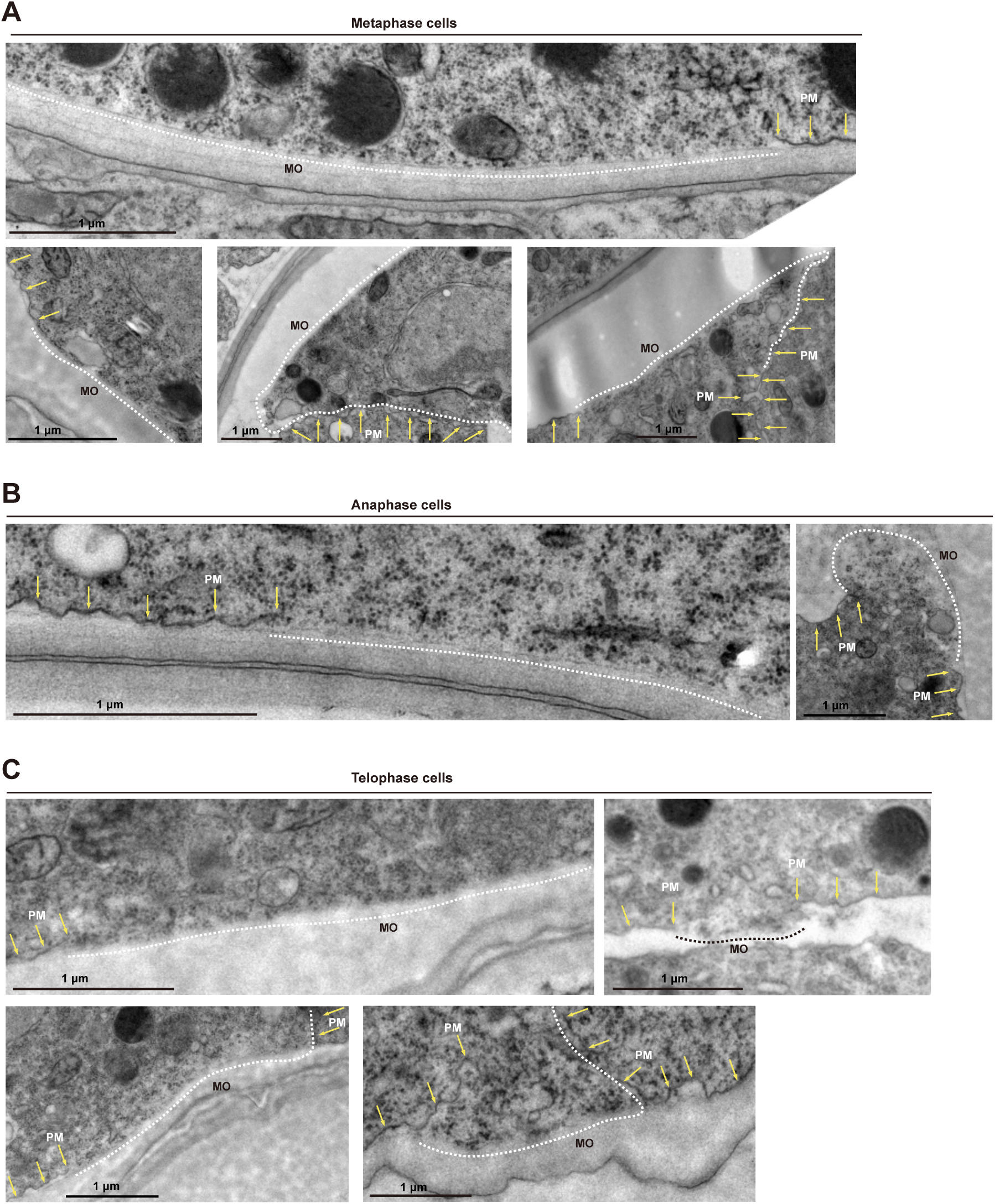
Membrane discontinuities in cells during mitosis. A-C, Membrane discontinuities or openings (MO, white or black dashed lines) in cells at metaphase (A), anaphase (B) or telophase (C). Yellow arrows points to plasma membranes (PMs).

**Figure S4.**
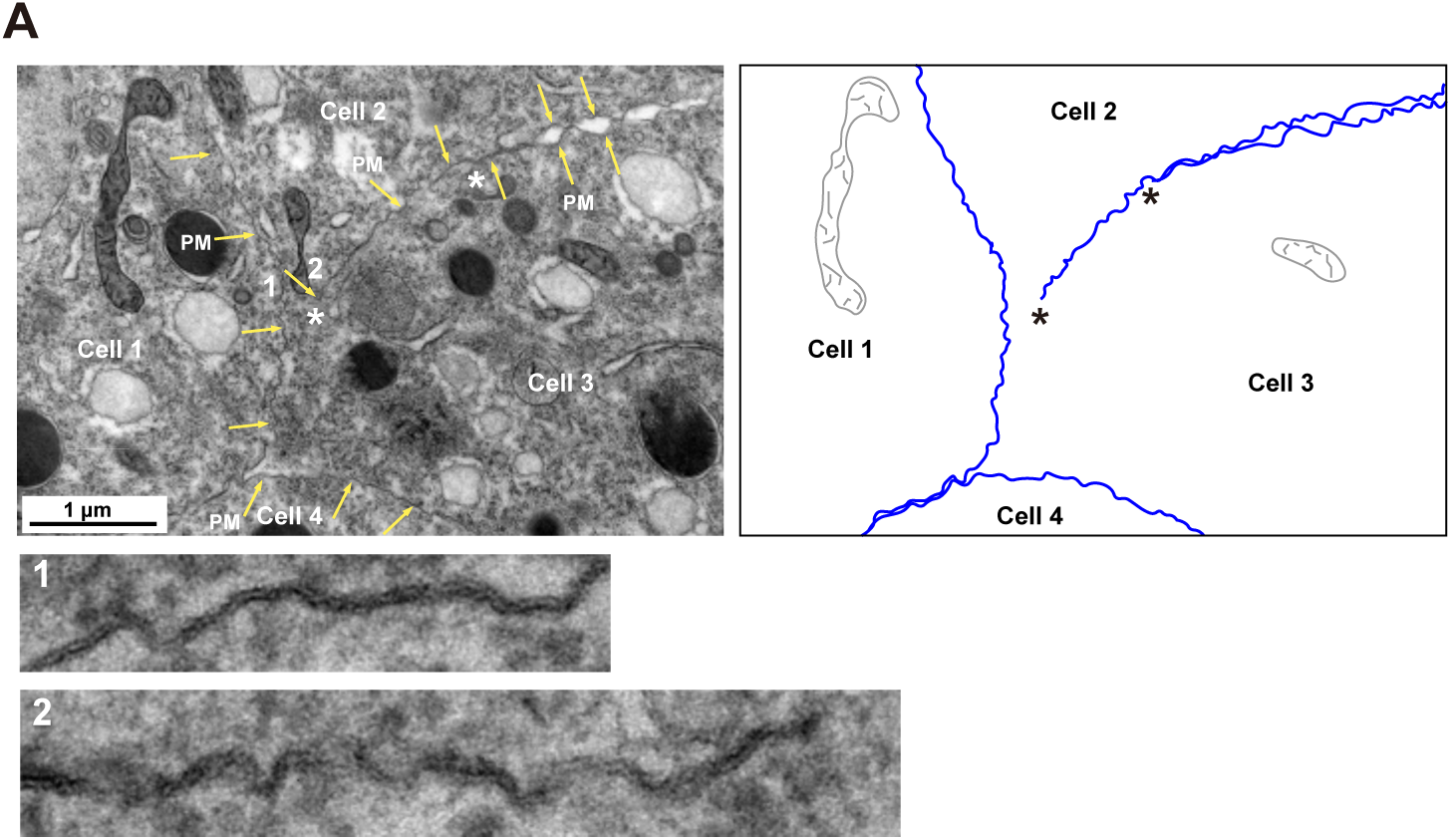
Cytosol-immersed single membranes. A, Cytosol-immersed single membranes separating the cytoplasmic regions of two adjacent cells. Enlarged views of regions 1 and 2 are shown below. Plasma membranes (PM) are indicated by yellow arrows or blue lines. Asterisks indicate membrane tips.

**Video S1. 3D reconstruction of an entire early *C. elegans* embryo.**

Membrane openings are highlighted in blue. Plasma membranes are in white.

**Video S2. 3D reconstruction of the metaphase cell in Figure 4B**.

The plasma membrane is shown in red; the condensed chromosome is in blue, and exposed cytoplasm is in green.

**Video S3. 3D reconstruction of the membrane opening on a tri-nucleated cell in Figure 4E**.

3D reconstruction of membrane ruptures and adjacent plasma membranes. Individual membranes, each associated with a distinct nucleus, are color-coded (purple, yellow, or blue).

**Video S4. Dynamic movement of membrane openings in Figure 7A**.

Green signal is GFP::PH(PLC1δ1).

**Video S4. Dynamic movements of nuclear division and membrane openings in Figure 7B**.

Green signal is GFP::PH(PLC1δ1). Red signal is mCherry::H2B.

